# Biological nitrification inhibition (BNI) in wheat for climate adaptation in acidic and alkaline soils

**DOI:** 10.1101/2025.11.06.686715

**Authors:** Adrián Bozal-Leorri, Carmen González-Murua, Iratxe Zarraonaindia, Daniel Marino, Cesar Arrese-Igor, Mª Begoña González-Moro

## Abstract

Anthropogenic disturbances to nitrogen (N) cycling, particularly through agricultural N use, have intensified nitrous oxide (N_2_O) emissions. Developing wheat lines with biological nitrification inhibition (BNI) is a promising strategy to reduce such emissions. However, the effectiveness of BNI depends not only on plant characteristics, but also on how they interact with environmental factors such as soil pH and elevated atmospheric CO_2_ (eCO_2_). This study evaluated the response of a BNI-capable wheat line (*Triticum aestivum* cv. ROELFS) to eCO_2_ under contrasting soil pH during the 30 days following ammonium fertilization. Two wheat lines (ROELFS-Control and ROELFS-BNI) were grown under ambient (400 ppm) and elevated (800 ppm) CO_2_ in acidic (pH 5.3) and alkaline (pH 8.8) soils. N_2_O emissions, nitrifying and denitrifying microbial communities, and soil chemical properties were monitored to assess plant–soil–microbe interactions. ROELFS-BNI consistently reduced N_2_O emissions under all conditions by reducing archaeal (Nitrososphaeraceae) and bacterial nitrifiers (Nitrosomonadaceae, Nitrospiraceae) without major shifts in overall microbial composition, indicating high specificity of BNI exudates. Elevated CO_2_ effects on N_2_O emissions were pH-dependent. In acidic soil, eCO_2_ increased emissions in ROELFS-Control but not in ROELFS-BNI, likely due to the enrichment of complete denitrifiers (e.g. Rhodanobacteraceae). However, in alkaline soil, eCO_2_ reduced N_2_O emissions in both Control and BNI lines, especially the latter, which was associated with a higher abundance of N_2_O-reducing denitrifiers (e.g. Burkholderiaceae). This study highlights the potential of ROELFS-BNI wheat as a sustainable practice to mitigate N pollution adaptable to diverse soils and predicted CO_2_ atmospheric conditions.

## 1. Introduction

Anthropogenic activities have significantly altered global biogeochemical cycles, with nitrogen (N) playing a central role in environmental change. The extensive use of N fertilizers to sustain agricultural productivity has led to unintended consequences, including soil degradation, water contamination, and greenhouse gas emissions (Fowler et al., 2013; Galloway et al., 2008). Among these, nitrous oxide (N_2_O) emissions are of particular concern due to their dual impact as a potent greenhouse gas and a major contributor to stratospheric ozone depletion (Ravishankara et al., 2009). Agriculture is responsible for approximately 49% of anthropogenic N_2_O emissions, mainly through microbial processes such as nitrification and denitrification in fertilized soils (Fowler et al., 2009; Li et al., 2016). Given that global food demand is projected to increase, the intensification of agricultural practices without sustainable management strategies will likely exacerbate N losses and their associated environmental impacts (Tilman et al., 2011). Thus, addressing agricultural N emissions is crucial for both mitigating climate change and ensuring long-term food security.

Soil and environmental factors play a crucial role in regulating N_2_O emissions from agricultural systems. Soil pH, in particular, is a key determinant of microbial processes involved in N transformations (Cameron et al., 2013). Nitrification is most efficient in soils with a pH between 6.5 and 8.0 (Sahrawat, 2008), and may be reduced in more acidic soils, potentially decreasing N_2_O production. Denitrification occurs across a wider pH range but is most efficient in neutral to slightly alkaline conditions (Šimek and Cooper, 2002). Nevertheless, in highly acidic soils, denitrification may be incomplete, which could increase the proportion of N_2_O emitted relative to the completely reduced molecular nitrogen (N_2_) (Wang et al., 2018). Additionally, climate change is expected to modify these dynamics. Elevated atmospheric CO_2_ (eCO_2_), for example, which is projected to double the current CO_2_ concentration (IPCC 2014), exceeding 800 ppm by the end of the century, can alter plant physiology and root exudation patterns (Shimono et al., 2019), potentially influencing N cycling and greenhouse gas fluxes (Andrews et al., 2019; Vega-Mas et al., 2023). Thus, increases in atmospheric CO_2_ can enhance plant biomass and root activity, leading to increased organic matter inputs into the soil, which in turn can modify microbial-mediated N processes (Nie et al., 2013). Moreover, increased photosynthesis and plant growth under eCO_2_ will demand enhanced N inputs by the plant to meet food quality. Moreover, under these conditions increased ammonium (NH_4_ ^+^) uptake over nitrate (NO_3_ ^-^) assimilation can be favoured by certain crops (Rubio-Asensio and Bloom, 2016), which offers an opportunity to enhance nitrogen use while reducing nitrogen emissions. Understanding how both interacting factors (soil pH and eCO_2_) influence nitrogen cycling is essential for developing targeted strategies to reduce environmental pollution in agricultural systems. Given their combined effects on nitrification and denitrification, it is plausible that they play a key role in shaping N_2_O emission patterns.

To mitigate N losses and improve N use efficiency (NUE), synthetic nitrification inhibitors (SNIs) have been widely used in agriculture over the past two decades. These compounds function by suppressing the activity of ammonia-oxidizing archaea (AOA) and bacteria (AOB), thereby slowing down the conversion of NH_4_ ^+^ to NO_3_ ^-^ in soils (Ruser and Schulz, 2015; Beeckman et al., 2018). However, their effectiveness varies depending on soil type and climatic conditions, and their adoption is often limited by economic and regulatory constraints (Subbarao et al. 2013). Biological nitrification inhibition (BNI) has emerged as a promising and sustainable approach to control nitrification and subsequent N_2_O emissions (Subbarao et al., 2015). Molecules with BNI capacity (Biological Nitrification Inhibitors, BNIs) are naturally released by certain plant species that selectively inhibit ammonia-oxidizing microorganisms in the soil, thereby reducing NO ^-^ accumulation and N_2_O emissions (Lu et al., 2024). This eco-friendly strategy has the potential to enhance soil health, minimize N losses, and decrease reliance on chemical inputs in agricultural systems.

Although the capacity for BNI was first reported in grasses, including the wild wheat relative *Leymus racemosus*, wheat has traditionally been considered a species that lacks BNI capacity in its root system (Subbarao et al., 2007). Wheat plays a fundamental role in global food security, serving as a primary source of carbohydrates and proteins for billions of people (Schnurbusch 2019; FAOSTAT 2022). Furthermore, the fact that many wheat agrosystems are managed under intensive fertilization due to the high N requirement, leads to the expectation that N_2_O emissions from N fertilization could reach up to 7.6 Tg N year^-1^ by 2030 (Reay et al., 2012; Tilman et al., 2002). The introgression of their BNI-controlling chromosome region from *L. racemosus* into elite high-yielding wheat cultivars has succeeded in the development of BNI-enhanced wheat lines (Subbarao et al., 2021). These elite lines hold significant potential for maintaining high productivity while mitigating N pollution, making them a valuable tool for sustainable agriculture (Subbarao and Searchinger, 2021). One such line, ROELFS-BNI, has shown the ability to reduce N_2_O emissions under controlled conditions compared to its parental line (Bozal-Leorri et al., 2022; Subbarao et al., 2021). However, its effectiveness on modulating N_2_O emissions under different scenarios, such as the combined influence of soil pH and eCO_2_, remains largely unexplored.

In this study, we hypothesized that ROELFS-BNI wheat plants would exhibit lower N_2_O emissions than their parental line across contrasting soil pH conditions and under elevated atmospheric CO_2_, primarily through the inhibition of nitrification-derived nitrogen losses, without compromising plant growth. Therefore, we evaluate the potential of the ROELFS-BNI wheat line to mitigate N pollution under future climate conditions, focusing on eCO_2_ and two soil pH levels, acidic and alkaline. This experiment was designed to address the existing gap in knowledge regarding the combined effects of soil pH and elevated CO_2_ on the effectiveness of BNI traits in reducing N_2_O emissions. By examining key microbial groups targeted by BNIs, we aim to uncover the mechanisms driving nitrogen transformations and greenhouse gas dynamics. We also assess how eCO_2_ influences BNI trait–microbe–soil interactions. This study advances the understanding of BNI technology as a tool for sustainable agriculture and contributes to the development of climate-resilient nitrogen management strategies that balance productivity with environmental stewardship.

## 2. Methods

### 2.1. Experimental setup and growth conditions

The experiment was carried out in microcosms in a growth chamber under controlled conditions with a daily regimen of a 14/10 h day/night cycle, an average day/night temperature of 23/18 ℃ and a day/night relative humidity of 60/70%. Two CO_2_ conditions were studied: ambient CO_2_ (aCO_2_) at 400 ppm and elevated CO_2_ (eCO_2_) at 800 ppm. Two genetic stocks of elite wheat (*Triticum aestivum*) from ROELFS variety were tested in this study: ROELFS-Control (the parental line) and ROELFS-BNI (Subbarao et al., 2021). For germination, seeds were placed in square Gosselin plates at 5 °C in darkness for 7 days. Subsequently, the seeds were transferred to trays containing a perlite:vermiculite mixture (1:3; v:v) and kept at 20 °C for 4 days.

### 2.2. Soil collection and preparation

Soil was collected from a 0-30 cm layer from fields in February 2023. Acidic soil (pH 5.3) was collected from a wheat field in Mabegondo (A Coruña, Spain) (43° 14’ N, 8° 16’ W, 700 m above sea level). Similarly, alkaline soil (pH 8.8) was collected from a barley field in Tajonar (Navarre, Spain) (42° 46’ N, 1° 36’ W, 584 m above sea level). Soil properties were determined in the analytical services of Neiker (Derio, Spain) and are shown in Table 1. Roots and stones were meticulously removed using a 5 mm sieve. The soils were air-dried, homogenized, and stored at 4°C until the commencement of the experiment. As part of the pot preparation, soils were mixed with sand at a 3:1 ratio (v:v) to improve porosity before the start of the experiment. The experimental design consisted of 24 pots (5 L volume, 20 cm diameter), with 12 pots filled with 5.5 kg of an acidic soil–sand mixture and 12 with 5.5 kg of an alkaline soil–sand mixture (hereafter referred to as soil). Six pots of each soil type (acidic and alkaline) were assigned to aCO_2_, and the remaining six of each soil type to eCO_2_. To stimulate the activity of microorganisms involved in the N cycle, soils in each pot were activated following the procedure described by Torralbo et al. (2017). Additionally, the soil was rehydrated with deionized water to achieve 60% water-filled pore space (WFPS). WFPS was calculated as in Linn and Doran (1984) following the equation:

**Table 1.**
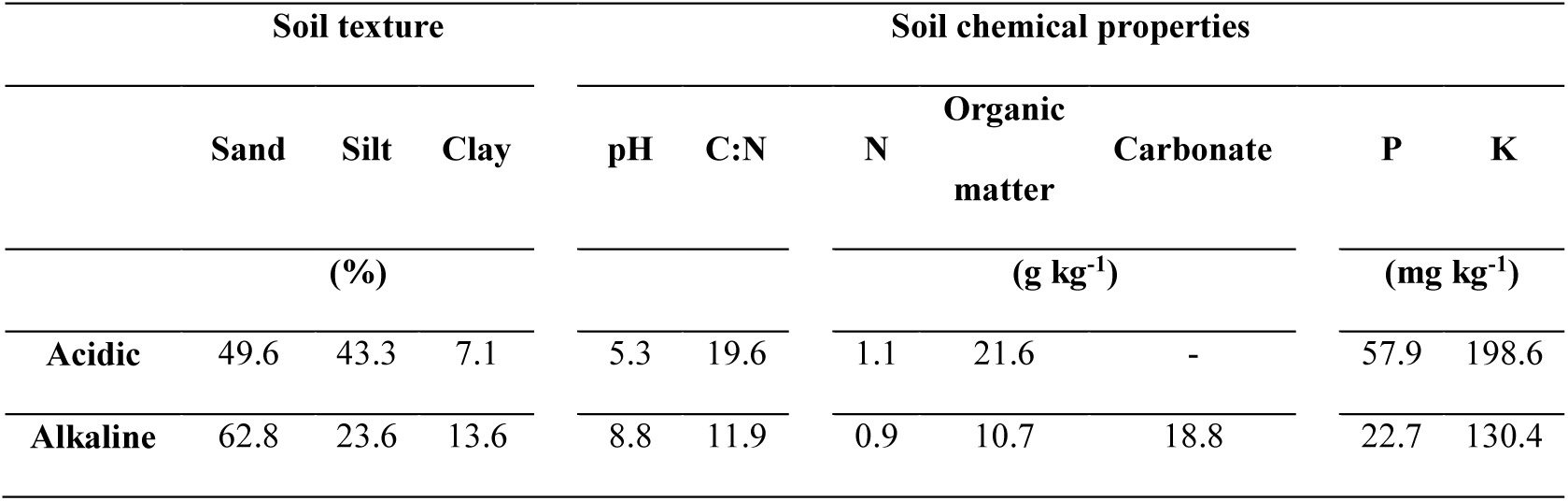
Physical and chemical properties of the acidic (43° 13’ N, 8° 16’ E, A Coruña, Galicia, Spain) and alkaline (42° 46’ N, 1° 37’ W, Tajonar, Navarre, Spain) soils collected in 0 – 30 cm depth layer after being mixed with sand.

WFPS = (soil gravimetric water content x bulk density) x (1 - (bulk density / particle density))-1

Particle density was assumed to be 2.65 Mg m^-3^ and soil bulk density was determined in the laboratory, resulting in a value of 1.27 Mg m^-3^ and 1.35 Mg m^-3^ for the acidic and alkaline soil, respectively. After 15 days of soil activation, pots from each soil pH level and CO_2_ concentration were divided into 2 groups (one for ROELF-Control line and other for ROELFS-BNI). This resulted in three biological replicates per treatment combination (2 soil pH × 2 CO_2_ × 2 wheat lines; n = 3), giving the abovementioned total of 24 pots. Six seedlings from the corresponding Control and BNI-lines were then placed in each pot, maintaining the soil at 60% of WFPS for an additional 15 days to allow plant establishment and growth. After, the experiment started with the addition of nitrogen in a single dose to the soil surface in an equivalent dose to 180 kg N ha^−1^, which was achieved by adding 2,665 mg of ammonium sulfate ((NH_4_)_2_SO_4_) in granular form. All pots were watered every two days to maintain 60% WFPS until the end of the experiment, which lasted for 30 days post-fertilization (DPF).

### 2.3. Aboveground biomass measurement

Aboveground biomass production of wheat was measured as dry weight (DW) per plant at 30 days post-fertilization (corresponding to 45 days post-transplanting). To do so, plants were dried at 80°C in a convection oven for 72 hours until a constant DW was achieved.

### 2.4. N_2_O and CH_4_ emission measurements

N_2_O and CH_4_ soil emissions were quantified employing the closed chamber method (Chadwick et al., 2014). On sampling days, chambers (20 cm diameter x 22 cm height) were inserted into the pots, reaching the edge of the soil, and subsequently removed after measurement. To ensure accurate measurements and subsequent data analysis, sampling was conducted following the protocols outlined in Hutchings et al. (2024) and the guidelines provided by de Klein et al. (2020). Briefly, sampling began after fertilization and was conducted three times per week for two weeks. Sampling was then reduced to twice per week for the following two weeks until N_2_O emissions stabilized (25 days). Gas samples were collected immediately after closing the chambers and again after 45 minutes. From each chamber, 35 mL of gas was extracted and stored at overpressure in pre-evacuated 20 mL glass vials. Throughout the experiment, the linearity of fluxes was regularly checked taking samples at 0, 15, 30, 45 and 60 min. Samples were analysed in a gas chromatograph (Agilent, 7890A) equipped with an electron capture detector for N_2_O detection and a flame ionization detector for CH_4_ determination. A capillary column (IA KRCIAES 6017:240 °C, 30 m × 320 µm) was used, and gas samples were injected utilizing a headspace auto-sampler (Teledyne Tekmar HT3). Standards of N_2_O and CH_4_ were analysed as controls.

### 2.5. Soil mineral nitrogen analysis

Soil mineral N content (NH_4_ ^+^-N and NO_3_ ^-^-N) was determined on day 30 post-fertilization (DPF). At the end of the experiment, each pot was destructively sampled, and its soil was thoroughly homogenized. A subsample of 100 g fresh soil was then taken and mixed with 1 M KCl (200 mL) and shaken at 165 rpm for one hour. The soil solution was filtered first through Whatman n°1 filter paper (GE Healthcare) and then through Sep-Pak Classic C18 Cartridges (125 Å-pore size; Waters), to remove particles and organic matter, respectively. The Berthelot method was followed to quantify the NH_4_ ^+^ content (Patton and Crouch, 1977), while the NO_3_ ^-^ content was determined according to Cawse (1967).

### 2.6. Quantification of microbial total abundance and community analysis

Bacterial and archaeal total abundance, as well as the abundance of functional marker genes related to nitrification and denitrification in the soil were quantified using quantitative polymerase chain reaction (qPCR) at the end of the experiment (30 DPF). DNA was extracted from the equivalent of 0.25 g dry soil taken from homogenized fresh soil samples (same material used for mineral N determination) using the PowerSoil DNA Isolation Kit (Qiagen), following the modifications outlined in Harter et al. (2014). Total bacterial and archaeal abundance (*16S rRNA*), genes associated with nitrification (bacterial and archaeal *amoA*) and denitrification (*nirK*, *nirS*, *nosZI* and *nosZII*) were amplified following the procedures described by Bozal-Leorri et al. (2022). The copy number of the target gene per gram of dry soil was calculated according to a modified equation described in Behrens et al. (2008): (number of target gene copies per reaction × volume of DNA extracted) / (volume of DNA used per reaction × gram of dry soil extracted). Additionally, archaeal and bacterial community diversity and composition were assessed through Illumina amplicon sequencing. The *16S rRNA* gene was amplified using the primer pairs Arch519/Arch915 (Herfort et al., 2009) for archaeal communities and 341F/805R for bacterial communities (Herlemann et al., 2011). Dual-indexed amplicons were generated using the Nextera XT Index Kit version 2. Libraries were purified using the CleanNGS kit (CleanNA, Waddinxveen, Netherlands) and sequencing was performed separately for each microbial fraction on an Illumina MiSeq platform at the Genotyping Service of the University of the Basque Country (SGiker) using MiSeq Reagent Kit version 3 (PE 2 × 300 bp, 600 cycles). Sequences were quality-filtered and demultiplexed using QIIME 2 (Bolyen et al., 2019), with denoising, merging, chimera removal, and amplicon sequence variant (ASV) determination performed via DADA2. Taxonomic classification of archaeal and bacterial sequences was conducted against Silva reference database using a prefiltered classifier based on Silva 138 99% OTUs full-length sequences.

### 2.7. Statistical analysis

Statistical evaluation of the data was carried out with QIIME2 and SPSS statistical software package (2016, IBM SPSS Statistics for Windows, Version 24.0. Armonk, NY, IBM Corp). Soil samples community richness and evenness were calculated through Chao1 and Shannon indices, respectively, rarifying the ASV table to 15,500 and 14,800 **s**equencing depth for bacteria and archaea, respectively. Mean comparison of all data was performed using two-way analysis of variance (Supplementary Table 2) with Duncan’s multiple range test for separation of means between different treatments. A p-value < 0.05 was considered to indicate statistically significant differences. Because amplicon sequence data are compositional in nature, relative abundances were transformed using the centered log-ratio (CLR) transformation prior to statistical analyses, following the recommendations of Gloor et al. (2017). The statistical tests were thus performed on CLR-transformed data, while untransformed relative abundances are presented in figures for clarity. This approach avoids the constraints inherent to compositional data and ensures the validity of subsequent ANOVA analyses. Community dissimilarity between samples was assessed using the Bray-Curtis index, and the ANOSIM test was used to determine whether microbial community composition changed significantly between the studied factors (wheat genetic lines, soil pH levels and CO_2_ condition).

## 3. Results

### 3.1. BNI trait maintains the growth of wheat plants

The introgression of the *Leymus racemosus* genomic region that confers the BNI trait into wheat maintains the growth of the new lines, as there were no significant differences in dry biomass between Control and BNI plants after 30 days of fertilization, regardless of pH level or CO_2_ condition (Supplementary Table 1). As expected, elevated CO_2_ condition similarly promoted the growth of ROELFS wheat lines, both the Control and BNI lines, which exhibited 16% and 9% higher dry biomass compared to aCO_2_ in acidic and alkaline soil, respectively.

### 3.2. The presence of ROELFS-BNI wheat plants reduce N_2_O emissions but increase CH_4_ emissions

The peak of N_2_O emissions occurred between days 3 and 7 post-fertilization (DPF) in both soil types (Supplementary Fig. 1A and B), but with much higher intensity in alkaline soil as reflected by the cumulative N_2_O emission values (Fig. 1A). Regarding atmospheric CO_2_ effect on cumulative emissions, eCO_2_ condition increased 69% the N_2_O emissions in acidic soil with ROELFS-Control wheat plants compared to aCO_2_ condition (Fig. 1A). However, the presence of BNI wheat lines were effective in maintaining low N_2_O emissions, regardless of the CO_2_ conditions with a 57% and 62% reduction in aCO_2_ and eCO_2_, respectively. In alkaline soil, the highest cumulative emissions were observed in soil with ROELFS-Control wheat plants under aCO_2_ conditions. In this case, eCO_2_ conditions reduced 30% the N_2_O emissions. Although less effective than in acidic soil, ROELFS-BNI wheat plants were also able to decrease N_2_O emissions in alkaline soil, accounting for a 26% and 33% reduction in aCO_2_ and eCO_2_, respectively. Daily methane (CH_4_) emissions remained constant throughout the entire experiment, without displaying any relevant peak (Supplementary Fig. 1C and D). The CO_2_ conditions had great influence on CH_4_ emissions. Indeed, CH_4_ was emitted under aCO_2_, whereas it was consumed under eCO_2_ (Fig. 1B). The presence of ROELFS-BNI affected CH_4_ emissions only under aCO_2_ conditions, leading to an increment of 54% in acidic soil and 131% in alkaline soil compared to ROELFS-Control line.

**Fig. 1.**
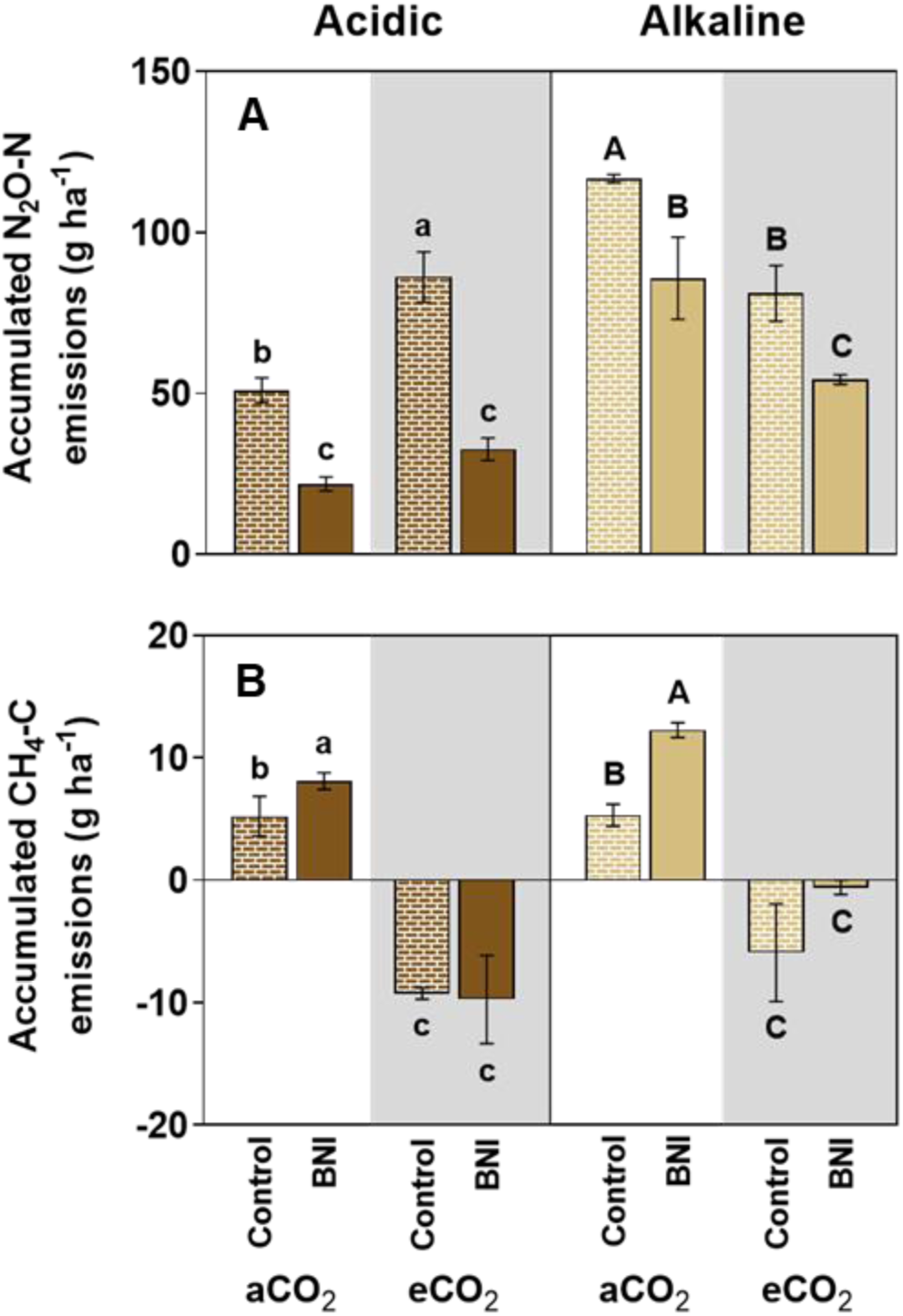
Cumulative N_2_O emissions (A) and CH_4_ emissions (B) during 25 days post-fertilization in acidic and alkaline soil under ambient (aCO_2_) and elevated (eCO_2_) CO_2_ conditions. Duncan’s post hoc test (p < 0.05) was used to denote significant differences with lowercase letters indicating differences in acidic soil and uppercase letters indicating differences in alkaline soil.

### 3.3. Nitrification and nitrifying microbial abundance are reduced by BNI-wheat

Regardless of the wheat genotype, more NH_4_ ^+^-N content was retained in acidic soil compared to alkaline soil (Supplementary Fig. 2A). No significant difference in NH_4_ ^+^-N retention was observed between ROELFS-Control and ROELFS-BNI wheat plants in acidic soil, regardless of CO_2_ concentration; however, NH_4_ ^+^-N content was lower under eCO_2_ (Supplementary Fig. 2A). In alkaline soil, ROELFS-BNI wheat plants exhibited slightly higher NH_4_ ^+^-N content compared to the Control lines only under eCO_2_ conditions. Regarding NO_3_^-^-N, there was no effect of BNI character under acidic soil and aCO_2_ conditions, whereas soils with ROELFS-BNI plants had lower NO_3_ ^-^-N content under eCO_2_ (Supplementary Fig. 2B). In alkaline soil under aCO_2_, ROELFS-BNI wheat plants showed higher NO_3_ ^-^-N content compared to ROELFS-BNI, while under eCO_2_ both wheat lines showed similar NO_3_ ^-^-N content.

As expected, both archaeal and bacterial community structures were significantly shaped by soil pH, samples clearly grouping among the axis based on this factor and ANOSIM (R 1.0 and *p* 0.001 for both archaea and bacteria). Consequently, all subsequent statistical analyses were stratified by soil type.

Soil cultivated with ROELFS-BNI wheat plants exhibited significantly greater archaeal richness and evenness compared to ROELFS-Control lines, irrespective of soil pH or CO_2_ conditions (Table 2A). However, despite these differences in diversity, ANOSIM analysis revealed no significant differences in the overall archaeal community composition between the two wheat lines.

**Table 2.** Archaeal (A) and bacterial (B) alpha diversity Chao1 and Shannon indexes in acidic and alkaline soil under ambient (aCO_2_) and elevated (eCO_2_) CO_2_ conditions after 30 days post-fertilization. Duncan’s post hoc test (p < 0.05) was used to denote significant differences with lowercase letters indicating differences in acidic soil and uppercase letters indicating differences in alkaline soil. Differences in beta diversity of archaeal community based on analysis of dissimilarity (ANOSIM) test (Bray-Curtis distances). *p*-values are based on 999 permutations.

| A |  | Archaea |  |  |  |  |  |
| --- | --- | --- | --- | --- | --- | --- | --- |
|  |  |  | Chao1 | Shannon |  | ANOSIM |  |
|  |  |  |  |  |  | R | *p*-value |
| Acidic | aCO<sub>2</sub> | Control | 59.5 ± 3.2 b | 3.16 ± 0.00 b | Control vs. BNI | 0.16 | 0.088 |
|  |  | BNI | 89.1 ± 4.3 a | 3.65 ± 0.12 a |  |  |  |
|  | eCO<sub>2</sub> | Control | 71.0 ± 1.2 b | 3.49 ± 0.17 b | Control vs. BNI | 0.20 | 0.071 |
|  |  | BNI | 93.3 ± 8.1 a | 3.68 ± 0.10 a |  |  |  |
| Alkaline |  | Control | 48.0 ± 2.5 B | 4.07 ± 0.02 B | Control vs. BNI | 0.11 | 0.063 |
|  | **aCO<sub>2</sub>** | **BNI** | 106.6 ± 13.5 A | 4.68 ± 0.11 A |  |  |  |
|  | **eCO<sub>2</sub>** | **Control** | 61.6 ± 3.5 B | 4.19 ± 0.04 B |  |  |  |
|  |  | **BNI** | 98.5 ± 6.9 A | 4.77 ± 0.08 A | **Control vs. BNI** | 0.12 | 0.055 |

| **B** |  | **Bacteria** |  |  |  | **ANOSIM** |  |
| --- | --- | --- | --- | --- | --- | --- | --- |
|  |  |  | **Chao1** | **Shannon** |  | **R** | **p-value** |
| **Acidic** | **aCO<sub>2</sub>** | **Control** | 1762 ± 95 a | 10.0 ± 0.0 a | **Control vs. BNI** | 0.03 | 0.331 |
|  |  | **BNI** | 1846 ± 10 a | 10.1 ± 0.1 a |  |  |  |
|  | **eCO<sub>2</sub>** | **Control** | 1206 ± 72 b | 9.6 ± 0.7 a | **Control vs. BNI** | 0.01 | 0.312 |
|  |  | **BNI** | 1135 ± 84 b | 9.6 ± 0.7 a |  |  |  |
| **Alkaline** | **aCO<sub>2</sub>** | **Control** | 1048 ± 182 B | 9.5 ± 0.2 B | **Control vs. BNI** | 0.00 | 0.450 |
|  |  | **BNI** | 1128 ± 118 B | 9.6 ± 0.1 B |  |  |  |
|  | **eCO<sub>2</sub>** | **Control** | 1603 ± 17 A | 10.1 ± 0.0 A | **Control vs. BNI** | 0.00 | 0.474 |
|  |  | **BNI** | 1782 ± 148 A | 10.1 ± 0.1 A |  |  |  |

The total abundance of archaea was not significantly affected by the BNI trait or CO_2_ conditions, with values ranging from 1 × 10^7^ to 2 × 10^7^ copies of *16S rRNA* g^-1^ dry soil (Supplementary Fig. 3A). The abundance of ammonia-oxidizing archaea (AOA) was significantly higher in acidic soil, exceeding 3 × 10^6^ copies of *amoA* g^-1^ of dry soil, while in alkaline soil, it remained below 1×10^6^ copies of *amoA* g^-1^ of dry soil (Fig. 2A). In acidic soil, the presence of BNIs reduced the abundance of AOA by 32% and 20% under aCO_2_ and eCO_2_ conditions, respectively. In alkaline soil, BNIs were also effective in reducing the abundance of AOA regardless of CO_2_ conditions, achieving reductions of 45% and 11% under aCO_2_ and eCO_2_ conditions, respectively.

**Fig. 2.**
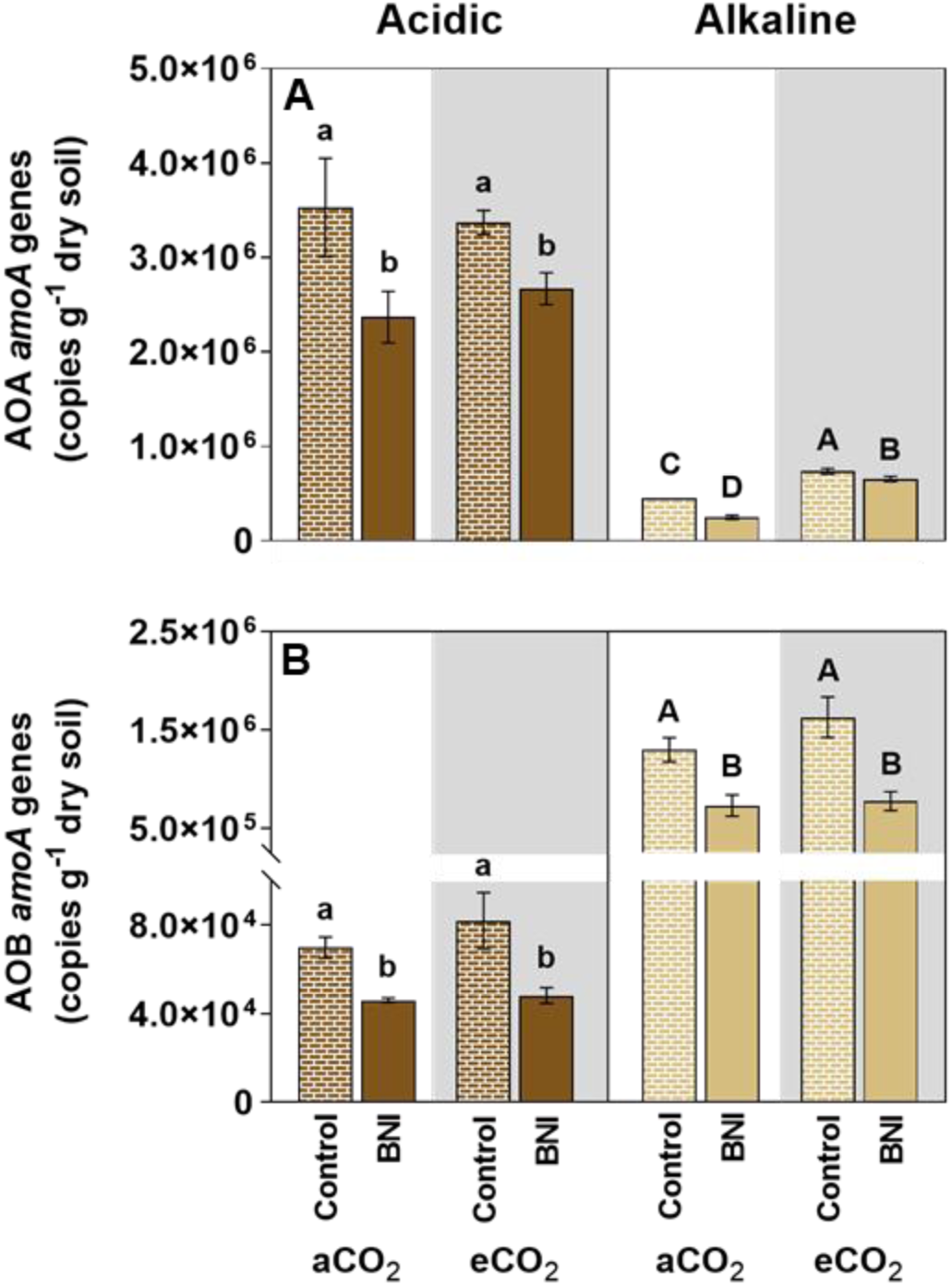
Total abundance of (A) ammonia-oxidizing archaea (AOA) and (B) ammonia-oxidizing bacteria (AOB) in acidic and alkaline soil under ambient (aCO_2_) and elevated (eCO_2_) CO_2_ conditions after 30 days post-fertilization. Duncan’s post hoc test (p < 0.05) was used to denote significant differences with lowercase letters indicating differences in acidic soil and uppercase letters indicating differences in alkaline soil.

The analysis of the archaeal population revealed that Nitrososphaeria was the dominant archaeal class under all conditions (Fig. 3A.1 and 3A.2), accounting for 81% to 95% of archaeal abundance. Other classes accounted for less than 1%, except for Thermoplasmata in acidic soils, where its relative abundance ranged from 2% to 14%. In both types of soils, the presence of BNIs reduced the relative abundance of Nitrososphaeria under both aCO_2_ and eCO_2_ conditions (Fig. 3A.1). In contrast, in acidic soils, the relative abundance of classes Thermoplasmata and Methanobacteria increased in soils with ROELFS-BNI plants, although for the latter this effect was observed exclusively under aCO_2_ conditions. Furthermore, in alkaline soils, Methanobacteria and Methanocellia classes showed increased relative abundance in soils with ROELFS-BNI plants; however, similar to acidic soils, this effect was observed only under aCO_2_ conditions. Within the Nitrososphaeria class, where the majority of ammonia-oxidizing archaea (AOA) are classified, the Nitrososphaeraceae family was the most abundant under all conditions, with a relative abundance ranging from 78% to 95% (Fig. 3B.1 and 3B.2). The presence of BNIs decreased the abundance of this family compared to soils with ROELFS-Control plants under both aCO_2_ and eCO_2_ conditions in both acidic (Fig. 3B.1 and 3C.1) and alkaline (Fig. 3B.2 and 3C.2) soils.

**Fig. 3.**
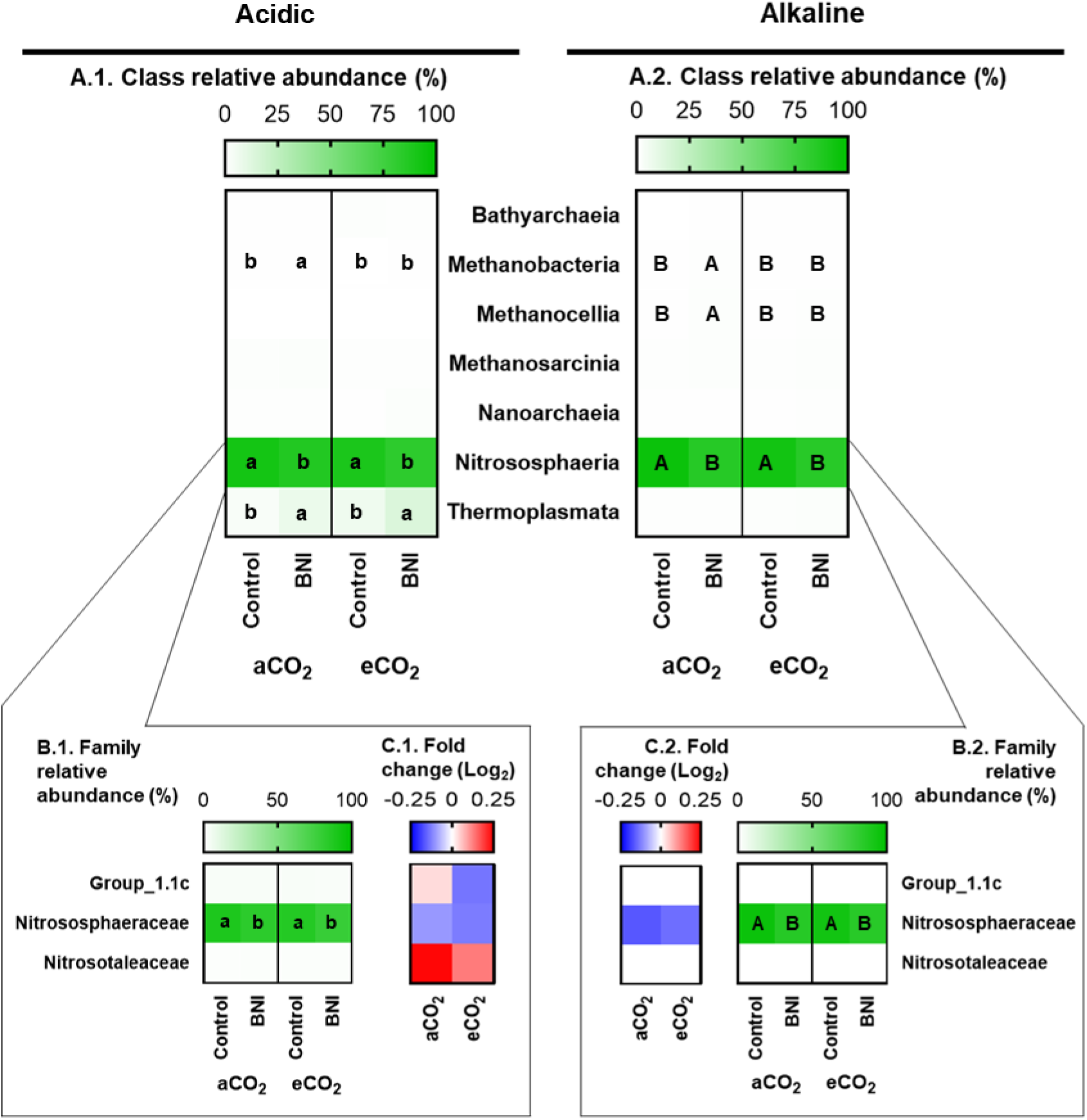
Relative abundance of archaeal classes (A.1 and A.2), relative abundance (B.1 and B.2) and variation in abundance of families from Nitrosphaeria class (C.1 and C.2) in soil with ROELFS-BNI wheat plants compared to soil with ROELFS-Control wheat plants. Data are presented for acidic soil (A.1, B.1, and C.1) and alkaline soil (A.2, B.2, and C.2). Statistical analyses were performed on CLR-transformed data and significant differences were determined using Duncan’s post hoc test (p < 0.05). Lowercase letters indicate significant abundnace differences in acidic soil, while uppercase letters indicate significant differences in alkaline soil.

Regarding bacteria, its total abundance was not significantly affected by the presence of the BNI trait, nor by CO_2_ condition with values ranging from 1.0 × 10^8^ to 1.5 × 10^8^ copies *16S rRNA* g^-1^ of dry soil (Supplementary Fig. 3B). Similarly, bacterial diversity and the overall composition in the soil was similar for soil cultivated with both genetic lines (Table 2B). In fact, bacterial richness and evenness were more significantly affected by CO_2_ conditions than by the presence of the BNI trait under both soil pH conditions (Table 2B). The impact of eCO_2_ had an opposite trend in acidic and alkaline soil. While eCO_2_ led to a decrease of Chao1 index in acidic soil, in alkaline soil an increase in Chao1 and Shannon indexes was observed (Table 2B), independent of the presence of the BNI trait.

The total abundance of ammonia-oxidizing bacteria (AOB) was significantly lower in acidic soil, remaining below 8 × 10^4^ copies *amoA* g^-1^ of dry soil, while in alkaline soil it exceeded 5 × 10^5^ copies *amoA* g^-1^ of dry soil (Fig. 2B). In both soils, BNIs effectively reduced the abundance of AOB under both CO_2_ conditions, achieving reductions of 34% at aCO_2_ and 41% at eCO_2_ for acidic soil and of 43% at aCO_2_ and 52% at eCO_2_ for alkaline soil.

Nitrifying bacteria accounted for no more than 3% of the relative bacterial abundance (Fig. 4). In acidic soil, among the AMO+HAO-containing bacteria, Mycobacteriaceae was the most abundant family (1.15%), followed by Nitrosomonadaceae, with relative abundances ranging from 0.46% to 1.12% (Fig. 4A.1). The relative abundance of AMO+HAO-containing bacteria was not affected by CO_2_ conditions. In ROELFS-BNI wheat plants, a decrease in the abundance of the Mycobacteriaceae, Nitrosomonadaceae, and Nitrospiraceae families was observed (Fig. 4A.1 and 4B.1) respect to ROELFS-Control wheat plants. In alkaline soil, Nitrosomonadaceae was the most prevalent family among the AMO+HAO-containing bacteria, with relative abundances ranging from 1.32% to 1.56% (Fig. 4A.2). The BNI trait in wheat reduced the relative abundance of the Nitrosomonadaceae and Nitrospiraceae families in the soil, but CO_2_ conditions did not affect the relative abundance of these nitrifiers (Fig. 4A.2 and 4B.2). Among HAO-containing bacteria, Isosphaeraceae was the most abundant family in acidic soil, but remained constant, regardless the level of atmosphreric CO_2_ and BNI trait. Similarly, in alkaline soils, neither the CO_2_ conditions nor the presence of the BNI trait affected the relative abundance of HAO-containing bacteria, with the exception of Pirellulaceae. This family, in addition to being the most abundant HAO-containing family, showed increased abundance under eCO_2_ conditions in alkaline soil where ROELFS-BNI wheat plants grew.

**Fig. 4.**
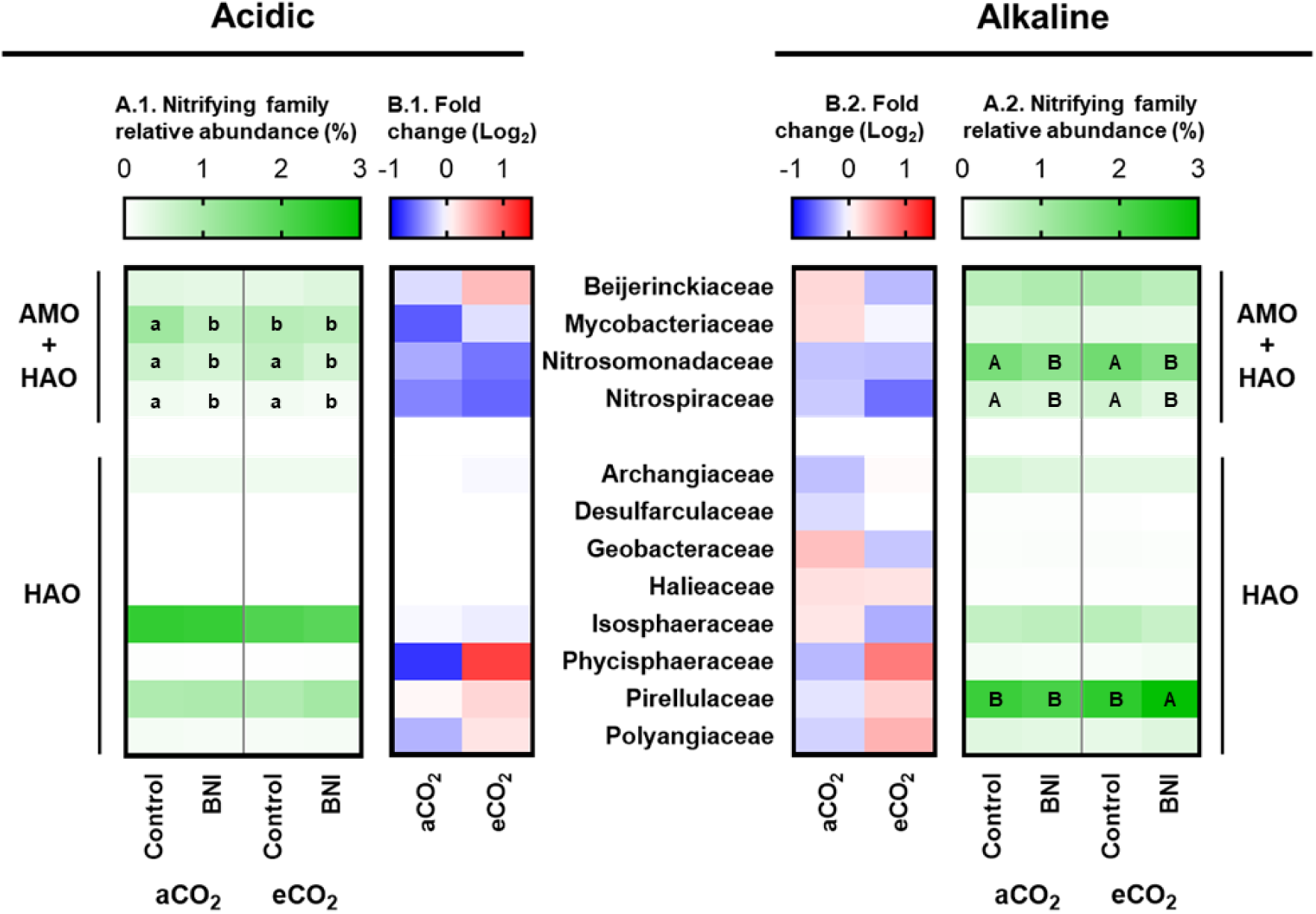
Relative abundance (A.1 and A.2) and variation in abundance (B.1 and B.2) of different nitrifying families in soil with ROELFS-BNI wheat plants compared to soil with ROELFS-Control wheat plants. Data are presented for acidic soil (A.1 and B.1) and alkaline soil (A.2 and B.2). Statistical analyses were performed on CLR-transformed data and significant differences were determined using Duncan’s post hoc test (p < 0.05). Lowercase letters indicate significant abundnace differences in acidic soil, while uppercase letters indicate significant differences in alkaline soil.

### 3.4. ROELFS-BNI wheat plants promote complete denitrifying bacteria abundance

The abundance of the *nirK* and *nirS* genes was lower in acidic soil than in alkaline soil (Fig. 5A and B). In acidic soil, ROELFS-BNI wheat lines reduced *nirK* abundance under eCO_2_ conditions, while in alkaline soil, the reduction occurred regardless of CO₂ levels. Similarly, BNIs significantly reduced *nirS* abundance in acidic soil by 39% and 57% under aCO_2_ and eCO_2_, respectively, but had no effect in alkaline soil. In contrast, *nosZI* abundance was similar in both soil types and increased by at least 60% in ROELFS-BNI lines under all conditions (Fig. 5C). Additionally, ROELFS-BNI enhanced *nosZII* abundance in alkaline soil (Fig. 5D).

**Fig. 5.**
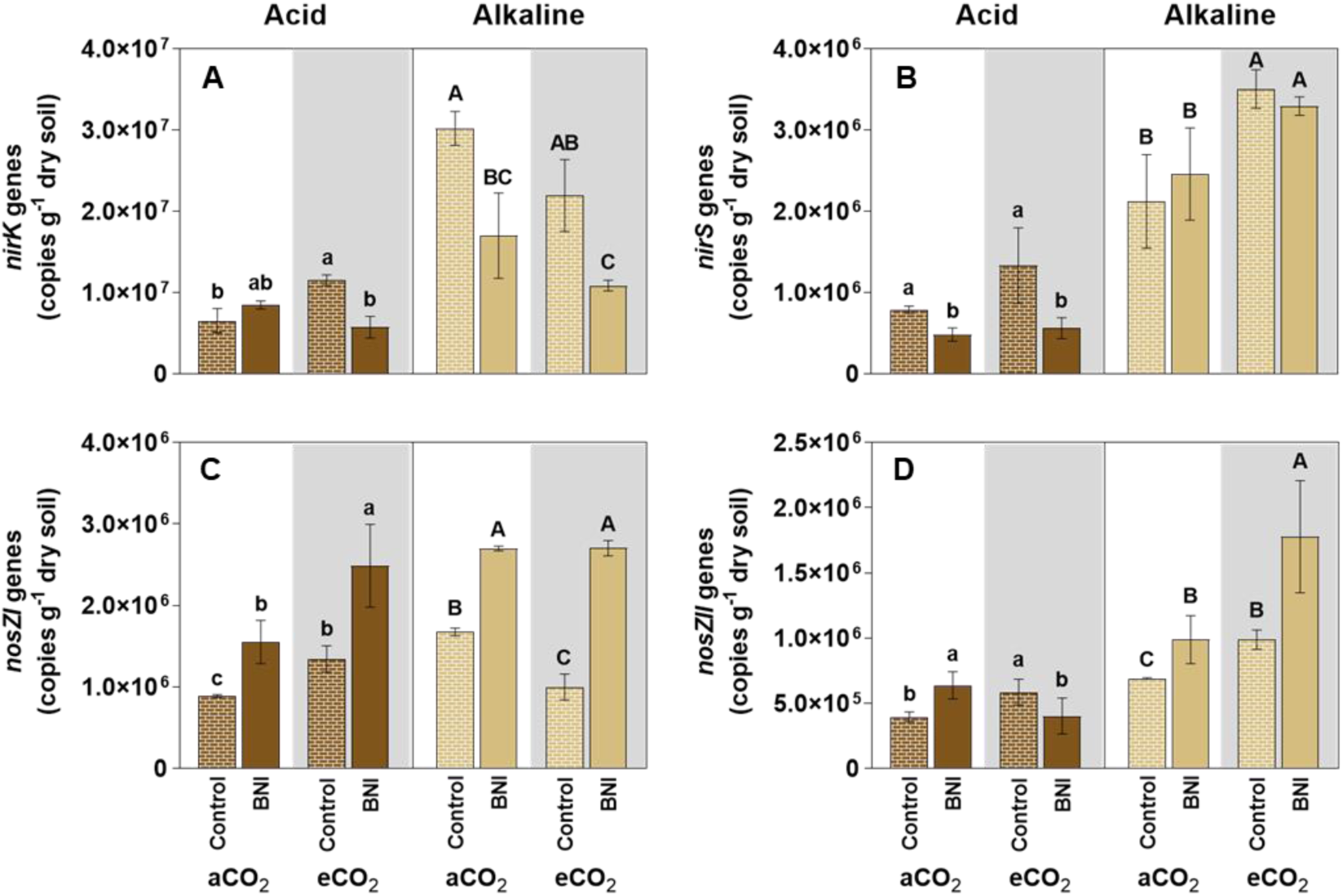
Total abundance of denitrifying bacteria containing (A) *nirK* (B) *nirS* (C) *nosZI* and (D) *nosZII* genes in acidic and alkaline soil under ambient (aCO_2_) and elevated (eCO_2_) CO_2_ conditions after 30 days post-fertilization. Duncan’s post hoc test (p < 0.05) was used to denote significant differences with lowercase letters indicating differences in acidic soil and uppercase letters indicating differences in alkaline soil.

Denitrifying bacterial families had relative abundances no higher than 3.1% (Fig. 6). In acidic soil, Gemmatimonadaceae (NO producers, 1.6% - 2.2%) and Burkholderiaceae (N_2_ producers, 2.3% - 3.1%) were dominant (Fig. 6A.1). CO_2_ conditions and BNIs had minimal effects, except for an increase in Gemmatimonadaceae and a decrease in Burkholderiaceae under eCO_2_. Rhodanobacteraceae (N_2_ producers) also increased with BNIs, particularly under aCO_2_. In alkaline soil, BNI-wheat had stronger effects. Burkholderiaceae (N_2_ producers, 1.7% - 2.9%) remained dominant, but other NO producers, including Anaerolineaceae, decreased under BNIs and aCO_2_. BNIs increased Herpetosiphonaceae under both CO_2_ conditions, while Micromonosporaceae and Pseudonocardiaceae declined, especially under eCO_2_ (Fig. 6A.2 and 6B.2). No effect was found among N_2_O producers. For N_2_ producers, Burkholderiaceae, Rhodanobacteraceae, and Flavobacteriaceae increased in BNI soils, with Flavobacteriaceae showing a stronger effect under eCO_2_.

**Fig. 6.**
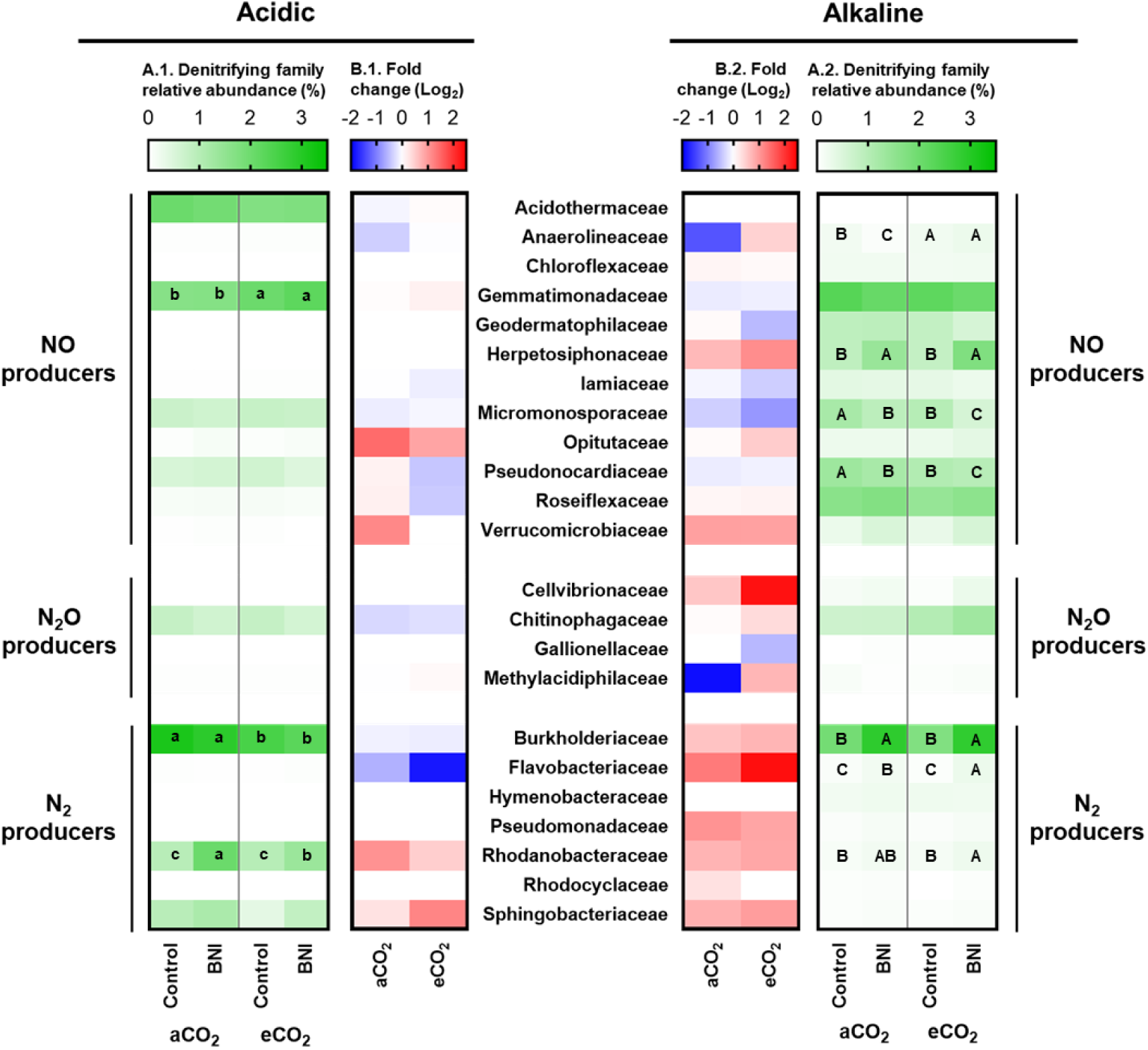
Relative abundance (A.1 and A.2) and variation in abundance of different denitrifying families (B.1 and B.2) in soil with ROELFS-BNI wheat plants compared to soil with ROELFS-Control wheat plants. Data are presented for acidic soil (A.1 and B.1) and alkaline soil (A.2 and B.2). Statistical analyses were performed on CLR-transformed data and significant differences were determined using Duncan’s post hoc test (p < 0.05). Lowercase letters indicate significant abundnace differences in acidic soil, while uppercase letters indicate significant differences in alkaline soil.

## 4. Discussion

### 4.1. ROELFS-BNI wheat line offers a promising solution to N emissions, even in the face of climate change challenges

BNI technology has been identified as a promising strategy for mitigating N losses (Subbarao et al., 2021). Here, we further demonstrated that BNI-ROELFS wheat plants are also effective in reducing the accumulated N_2_O emissions, therefore, mitigating pollution from intensive fertilization (Fig. 1A), under a predicted scenario of elevated CO_2_. As previously stated, N₂O is produced through the processes of nitrification and denitrification (Li et al., 2016). Although BNIs primarily reduce pollution by directly interfering with the nitrification process and its associated N₂O emissions (Prosser et al., 2020), the subsequent reduction in NO₃⁻ formation could also influence denitrification, which is the main pathway for N₂O production (Li et al., 2016). Therefore, a comprehensive understanding of the effects of BNIs on soil microbial groups involved in these two pathways, as well as considering different soil types (particularly varying pH levels) and climate-related factors, is crucial for developing sustainable agricultural practices that could be effective under a wide range scenario.

### 4.2. BNI-wheat is effective in reducing both archaeal and bacterial nitrifying population regardless of soil pH and atmospheric CO_2_ concentration

The use of the recently developed BNI-wheat lines is projected to reduce nitrification by up to 40% and enhance NUE by 16% by 2050 (Leon et al., 2022). BNIs typically show greater efficiency in acidic soils (Zhu et al., 2012), particularly affecting AOA (Kaur-Bhambra et al., 2022; Sarr et al., 2020), which are generally more abundant in these soil type (Hu et al., 2014). Nonetheless, archaea are also present in alkaline soils, making it essential to assess the effects of BNIs on these populations in such environments. To fill this knowledge gap, the effects of BNI-wheat on archaea community structure upon varying pH, and under ambient and elevated CO_2,_ were concurrently examined.

As expected, a higher relative abundance of AOA was observed in acidic soils. Importantly, BNIs effectively reduced AOA abundance not only in acidic soils but also in alkaline ones (Fig. 2A). Amplicon sequencing results further revealed that BNIs influenced the abundance of other archaeal groups and increased their richness in both soil types (Table 2A). Despite this, the overall composition remained stable, indicating that BNIs do not significantly alter the general structure of the archaeal community. Thus, the presence of BNIs in these new wheat lines would not compromise other vital soil ecosystem processes regardless of soil pH. Interestingly, in both acidic and alkaline soils, the relative abundance of the archaeal class Nitrososphaeria exceeded 81%, corroborating the findings of Saghaï et al. (2022) who also found it to be the dominant archaeal class in arable soils. This class includes all known AOA (Alves et al., 2018). Furthermore, within Nitrososphaeria, ROELFS-BNI wheat plants reduced the relative abundance of Nitrososphaeraceae, the most prevalent family (Fig. 3B.1 and 3B.2). Notably, the Group 1.1c, believed to be a non-ammonia oxidizing lineage due to the absence of *amoA* homologs (Weber et al., 2015), remained unaffected by BNIs. Importantly, this result evidences that BNIs specifically target AOA groups. Furthermore, AOA decrease was consistent across the varying pH levels and CO_2_ concentrations. Consequently, these findings suggest that BNI technology could be a viable strategy for mitigating nitrogen fertilizer pollution even in the face of climate change.

Aside from these global patterns, the abundance of particular groups were modified under certain conditions. For instance, the class Thermoplasmata increased only in acidic soils with ROELFS-BNI wheat plants (Fig. 3A.1). This class constitutes 5% of the archaeal abundance in global soil samples (Baker and Banfield, 2003) and ranged from 6% to 14% in our experiment. However, their ecology in agricultural soils remains largely uncharted, as Thermoplasmata have mainly been found in acidic mine drainage (Auguet et al., 2010). Hence, more efforts should focus on disentangling the functional potential of this group, together with the factors affecting its abundance and activity, since it is posited that they might perform metabolic functions such as methanogenesis, sulfur cycling, and even N_2_ fixation (Baker et al., 2020). The inhibition of nitrifying archaea through BNI release might have boosted this archaeal class independently of atmospheric CO_2_, which could have positive implications for the biogeochemical cycling of other important soil elements in acidic soils.

Together with AOA, AOB are also key players in the nitrification process in soils. The abundance of both groups is primarily influenced by soil pH and NH_4_ ^+^ availability. AOA typically dominate under acidic pH and low NH_4_ ^+^ concentrations, whereas AOB thrive under neutral to alkaline pH and higher NH_4_ ^+^ availability (Hink et al., 2018). Our findings confirm this general pattern, showing a higher abundance of AOB in alkaline soil compared to acidic soil (Fig. 2B). However, numerical dominance does not necessarily equate to functional dominance in nitrification (Ouyang et al., 2016). Recent studies suggest that AOB can be functionally active and may even be more important than AOA in acidic soils (Dai et al. 2018; Lin et al. 2018, 2021; Rojas-Pinzon et al. 2024). Indeed, using 1-octyne as an AOB inhibitor, Ouyang et al. (2017) reported that octyne-resistant AOA achieved maximum nitrification activity within an NH_4_ ^+^ concentration range of 0.005-0.02 mM. In our study, NH_4_ ^+^ concentration in acidic soils ranged from 40 mM NH_4_ ^+^ (138 mg NH4^+^ kg^-1^ dry soil) at the outset to about 0.25 mM NH_4_ ^+^ (0.86 mg NH_4_ ^+^ kg^-1^ dry soil) by the end of the experiment (Supplementary Fig. 2A). These values are substantially higher than those favouring AOA activity according to Ouyang et al. (2017), leading us to hypothesize that AOB were the main contributors to nitrification in our acidic soils, despite the expected ecological preference of AOA for such conditions.

The AOB abundance was not affected by eCO_2_ (Fig. 2B; Fig. 4A), corroborating previous studies (Barnard et al. 2004; Séneca et al. 2020; Bozal-Leorri et al. 2021). However, the BNI trait effectively reduced AOB abundance across both CO_2_ scenarios and soil types (Fig. 2B). This decline can be partly attributed to a decrease in the Nitrosomonadaceae family (Fig. 4), the predominant AOB in soil (Clark et al., 2021), as well as the Nitrospiraceae family (Fig. 4), which is thought to include complete ammonia-oxidizing bacteria (comammox) (Daims et al., 2015; Van Kessel et al., 2015). Interestingly, BNIs from ROELFS-BNI wheat lines specifically targeted bacterial families containing both the AMO and HAO enzymes, showing no impact on those with only the HAO enzyme (Fig. 4). Characterizing and identifying the chemical composition of BNI compounds in root exudates and their target enzymes is essential but poses significant challenges. To date, some molecules have been identified from various plants and crops, including *Brachiaria humidicola*, sorghum, rice, and maize (Subbarao et al. 2009; 2013; Lu et al. 2022; Otaka et al. 2022). Some BNI molecules specifically target the AMO enzyme, such as methyl 3-(4-hydroxyphenyl) propionate (MHPP) from sorghum and syringic acid from rice (Lu et al. 2022). Others, such as brachialactone from *B. humidicola* and sorgoleone from sorghum, inhibit the HAO enzyme (Subbarao et al., 2015). Unfortunately, the BNI molecules produced by *Leymus racemosus*, and consequently those released by BNI wheat lines, remain uncharacterized. Our findings clearly suggest that the root exudates from ROELFS-BNI may selectively inhibit the AMO enzyme while leaving the HAO enzyme unaffected (Fig. 4A.1 and 4A.2). Future research, particularly using untargeted metabolomics on root exudates from BNI wheat, will be crucial for discovering and understanding BNI compounds and their roles in soil physiology and health to support the implementation of sustainable agricultural practices.

### 4.3. The increase of total denitrifying bacteria also contributes to mitigation of N_2_O emissions

The impact of eCO_2_ on soil N_2_O emissions remains debated, as studies report increases, reductions, or no effect across different ecosystems (Wang et al., 2021). eCO_2_ influences key drivers associated with N_2_O emissions, such as soil carbon input, mineral N availability, moisture and related functional microbes (Liu et al., 2018). Therefore, we could expect that soils with contrasting pH respond differently to eCO_2_. In acidic soil, accumulated N_2_O emissions in presence of ROELFS-Control wheat plants increased under eCO_2_ conditions (Fig. 1A), aligning with the meta-analysis by Wang et al. (2021) who found that eCO_2_ generally promotes N_2_O emissions. The enhancement of N_2_O emissions due to CO_2_ levels in Control conditions could be linked to the rise of two important microbial families: the Gemmatimonadaceae family, which includes partial denitrifiers containing the *nirK* gene (DeBruyn et al., 2011), and a decreased abundance of Burkholderiaceae family, known for being complete denitrifiers (Zhu et al., 2018) (Fig. 6A.1 and B.1). Thus, this pattern could result in an increase in the production of N_2_O and, at the same time, a decrease in the overall reduction of such N gases to N_2_. On the other hand, the reduction in soil N_2_O emissions observed at both CO_2_ concentrations (Fig. 1A) mediated by ROELFS-BNI can be attributed not only to the decrease in AOA and AOB populations (Fig. 2A and B), but also to the positive effect of BNI trait on the Rhodanobacteraceae family (Fig. 6A.1 and B.1). Members of this family are complete denitrifiers, capable of reducing N_2_O at pH levels between 4 and 5.7 (Lycus et al. 2017; Hetz and Horn 2021), which falls within the acidic pH range of this experiment. Therefore, the BNI-driven increase on Rhodanobacteraceae may have played a crucial role in mitigating the expected rise in N_2_O emissions caused by elevated CO_2_ in acidic soils.

Unlike in acidic soils, eCO_2_ caused a decline in the accumulated N_2_O emissions in alkaline soil (Fig. 1A). This finding is consistent with a prior study in which a reduction in N₂O emission under eCO_2_ was also observed in soils of comparable alkaline pH where *Hordeum vulgare* was grown fertilized with SNIs (Bozal-Leorri et al., 2021). The analysis of microbial families revealed that eCO_2_ decreased the relative abundance of the families Micromonosporaceae and Pseudonocardiaceae (Fig 6A.2 and B.2), partial denitrifiers containing the nitrite reductase (NIR) enzyme, encoded by the *nirK* gene (Zumft, 1997). Dong et al. (2020) and Bozal-Leorri et al. (2021), demonstrated that the *nirK* gene decreased under eCO_2_ conditions in alkaline soils, a change they attributed to an increase in diverse microorganisms competing with denitrifiers. The significantly higher richness and evenness of bacterial communities observed under eCO_2_ compared to aCO_2_ (Table 2B), combined with the reduction in total *nirK* abundance reported in this study (Fig. 5A), further supports this hypothesis. Concerning the performance of BNI trait in alkaline soil, the presence of BNI probed to be effective in reducing the N_2_O emissions, with reductions of about 30% (Fig. 1A) relative to ROELFS-Control, under both CO_2_ conditions. This decline is likely linked to i) a reduction in nitrifying populations (AOA and AOB) (Fig. 2A and B) and ii) a decreased abundance of NO-producing bacteria families, including Micromonosporaceae and Pseudonocardiaceae (Fig. 6A.2 and B.2), alongside with an increased presence of complete denitrifying bacteria like Burkholderiaceae, Flavobacteriaceae, and Rhodanobacteraceae (Fig. 6A.2 and B.2). Although the increase in the latter 2 families was statistically significant, their overall relative abundances were low in the community (less than 0.1%). In addition, Rhodanobacteraceae is known to be unable to convert N_2_O to N_2_ at pH levels above 5.7 (Hetz and Horn, 2021), therefore, suggesting its limited role in denitrification under these conditions. In contrast, the Burkholderiaceae family was identified as the most abundant group among denitrifying bacteria, accounting for more than 3% of the total bacterial community. This family harbours the N_2_O-reducing gene (*nosZ*) (Zhu et al., 2018); indicating that it is crucial for the complete reduction of N_2_O to N_2_ in this particular soil. Indeed, in alkaline soils, synthetic nitrification inhibitors (SNIs) based on dimethylpyrazole have been found to increase the abundance of the *nosZ* genes (Barrena et al. 2017; Torralbo et al. 2017; Friedl et al. 2020; Bachtsevani et al. 2021; Huérfano et al. 2022). One postulated hypothesis is that the inhibition of AOB growth will free the copper (Cu) necessary for the AMO enzyme (Gilch et al., 2010) to be used by bacteria containing the N_2_O-reducing enzyme (Corrochano-Monsalve et al., 2021), which also requires Cu as a cofactor (Glass and Orphan, 2012). Similarly, BNI diminishes AOB populations, potentially increasing the availability of Cu for denitrifying bacteria. This mechanism might explain the observed increase in *nosZI* and *nosZII* gene abundances reported in this study (Fig. 5C and D). Nevertheless, additional research is necessary to elucidate the mechanisms underlying the potential impact of BNI on denitrifying bacterial families. This could entail tracking Cu availability in BNI-treated soils using isotopic labelling to assess whether Cu released from inhibited AOB activity is absorbed by N_2_O-reducing bacteria.

### 4.4. ROELFS-BNI wheat line does not affect methanotrophs but could indirectly influence methanogenic microorganisms

The presence of plants displaying BNI trait may not only influence the target microorganisms but could also produce off-target effects. Notably, since methane monooxygenase (MMO) enzyme shares a high degree of similarity with the AMO enzyme (Bédard and Knowles, 1989), it is plausible that BNIs influence methanotrophic microorganisms. However, this study detected only the Beijerinckiaceae family among methanotrophs (Fig. 4) (Tamas et al., 2014), with no significant differences found between the soils from the Control and those from BNI-wheat. This result further supports a highly selective action of BNIs against AMO-containing microorganisms. In contrast, the analysis of methanogenic microorganisms revealed an increase in the relative abundances of the archaeal classes Methanobacteria and Methanocellia in soils containing ROELFS-BNI wheat (Fig. 3A), which coincided with an increase in CH_4_ emissions (Fig. 1B). However, the finding that this effect was only observed under aCO_2_ conditions, and not under eCO_2_, suggests that it might not be a direct effect of the BNI trait but rather an indirect effect. For example, the reduction in the predominant archaeal class Nitrososphaeria (Fig. 3) in soils with ROELFS-BNI wheat lines might have allowed other archaeal classes to thrive and increase their abundance. Nevertheless, further metabarcoding studies under field conditions are needed to robustly confirm this hypothesis.

## 5. Conclusions

The use of BNI wheat holds strong potential for mitigating N_2_O emissions, as it inhibits nitrification and promotes conditions that limit emissions from denitrification. Our findings show that BNIs did not affect plant growth and effectively reduced N₂O emissions by influencing both microbial pathways. ROELFS-BNI wheat lines reduced nitrification by targeting AOA and AOB without significantly altering the overall composition of the soil archaeal and bacterial communities. In acidic soils, BNIs significantly reduced the abundance of the AOA containing *amoA* gene acting on Nitrososphaeraceae, the most abundant AOA family. In alkaline soils, BNIs effectively reduced AOB abundance acting on bacterial families such as Nitrosomonadaceae and Nitrospiraceae. The effective inhibition of nitrifying populations in both soil types regardless of the CO_2_ concentration reinforces the potential of this technology to reduce N losses in agricultural soils. Additionally, the *L. racemosus*-derived BNIs present in ROELFS-BNI wheat lines appear to specifically inhibit the AMO enzyme leaving HAO unaffected. The impact of eCO_2_ on N_2_O emissions varied with soil pH: it increased emissions in acidic soils under ROELFS-Control but not under ROELFS-BNI, likely due to shifts in denitrifier composition. In alkaline soils, eCO_2_ decreased N_2_O emissions overall, with further reductions in ROELFS-BNI treatments linked to enhanced abundance of complete denitrifiers such as Burkholderiaceae. Additionally, ROELFS-BNI wheat may alter methanogenic communities without affecting methanotrophic microorganisms, potentially influencing CH_4_ emissions. However, this effect was only detected under aCO_2_ conditions. To our knowledge, this study is the first to simultaneously investigate the effects of BNIs across two soils with differing pH levels, demonstrating BNI wheat lines effectively reduce both N_2_O emissions and nitrification in acidic and alkaline soils. These findings highlight the potential of BNI technology as a viable strategy to mitigate pollution from intensive fertilization. Moreover, they underscore the adaptability of this technology as a sustainable agricultural practice under future environmental conditions, particularly in response to rising CO_2_ levels driven by climate change.

## Abbreviations

AMO: Ammonia monooxygenase
AOA: Ammonia-oxidizing archaea
AOB: Ammonia-oxidizing bacteria
BNI: Biological nitrification inhibition
BNIs: Biological nitrification inhibitors
HAO: Hydroxylamine oxidoreductase
MMO: Methane monooxygenase
SNIs: Synthetic nitrification inhibitors

## Declarations

Ethics approval and consent to participate

Not applicable.

## Consent for publication

Not applicable.

Availability of data and material

The datasets supporting the conclusions of this article are available in the Qiita repository, under study ID 15753 (https://qiita.ucsd.edu/study/description/15753). Any further data will be available from the corresponding authors upon reasonable request.

## Author’s contribution

AB-L: Conceptualization, Data curation, Formal analysis, Investigation, Visualization, Writing – original draft. CG-M: Conceptualization, Supervision, Writing – review and editing. IZ: Data curation, Formal analysis, Writing – review and editing. DM: Funding acquisition, Investigation, Validation, Visualization, Writing – review and editing. CA-I: Conceptualization, Supervision, Validation, Writing – review and editing. MBG-M: Conceptualization, Funding acquisition, Investigation, Project administration, Supervision, Validation, Writing – review and editing.

## Acknowledgments

The authors thank for the technical and human support provided by SGIker, Phytotron Service of UPV/EHU and its European funding (ERDF and ESF).

