## Supplementary Material for "Biological nitrification inhibition (BNI) in wheat for climate adaptation in acidic and alkaline soils"

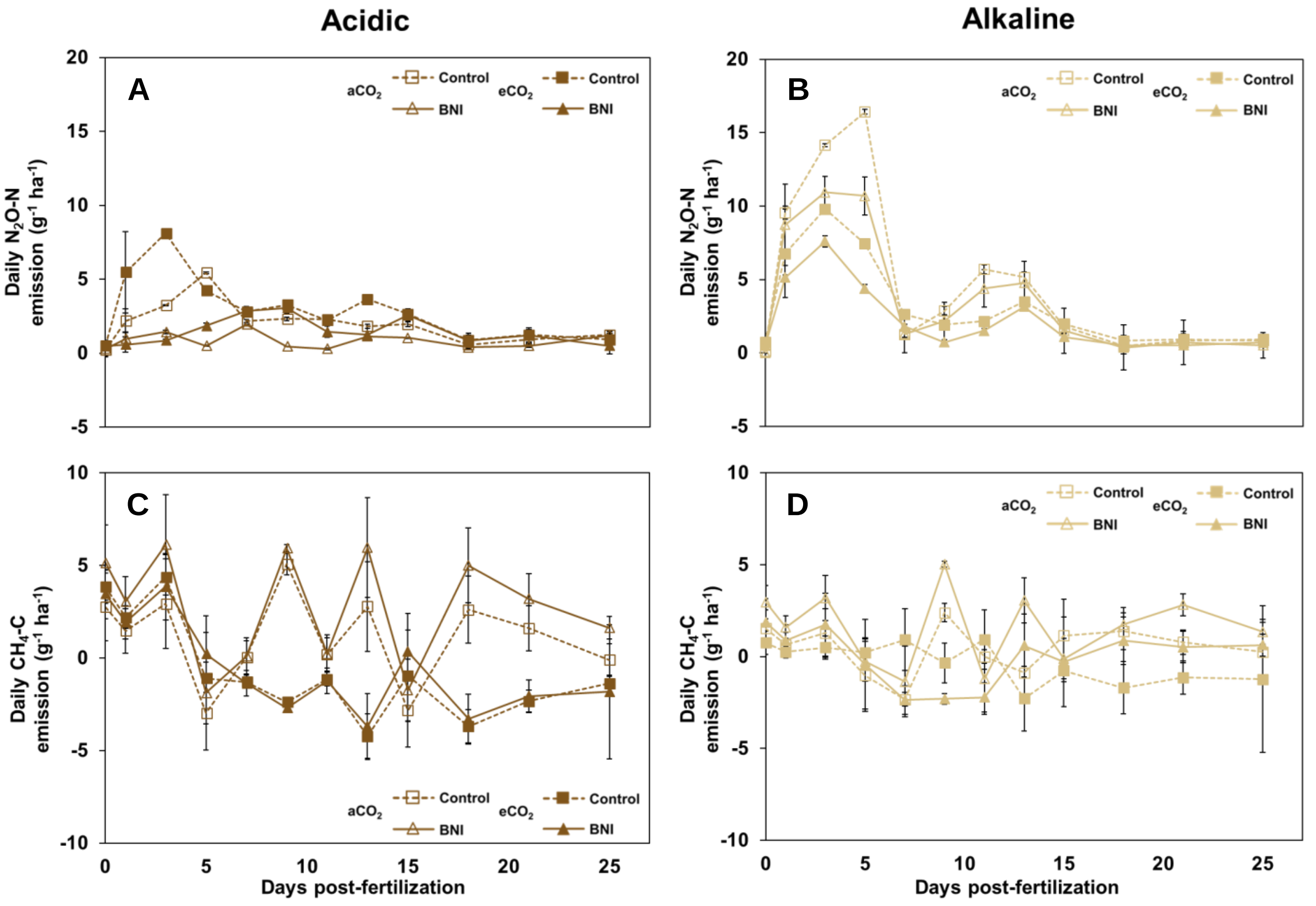


**Supplementary fig. 1.** Daily $N_2O$ emissions (A and B) and $CH_4$ emissions (C and D) during 25 days post-fertiliztion in acidic (A and C) and alkaline (B and D) soils under ambient ($aCO_2$) and elevated ($eCO_2$) $CO_2$ conditions.

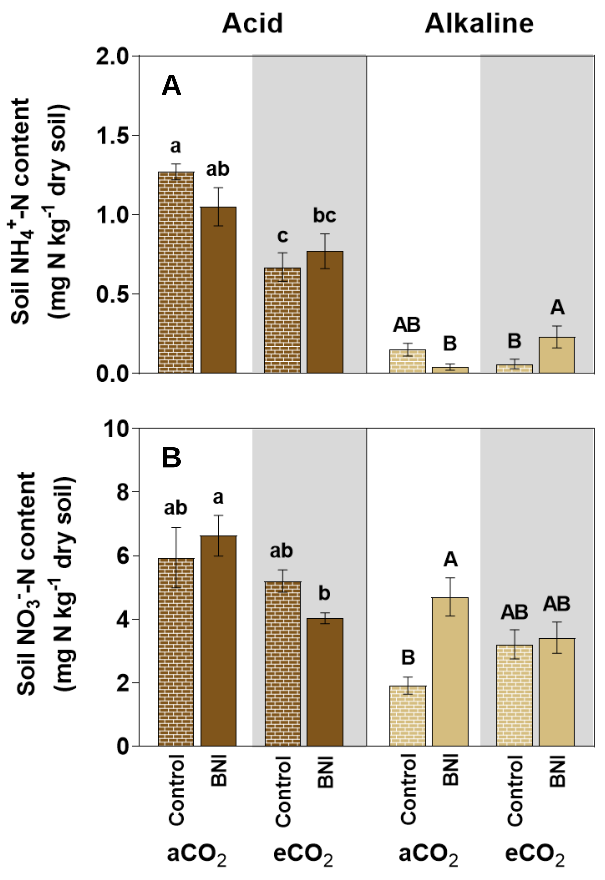


**Supplementary fig. 2.** Soil mineral nitrogen in form of $NH_4^+$ (A) and $NO_3^-$ (B) in acidic and alkaline soil under ambient ($aCO_2$) and elevated ($eCO_2$) $CO_2$ conditions after 30 days post-fertilization. Duncan's post hoc test ($p < 0.05$) was used to denote significant differences with lowercase letters indicating differences in acidic soil and uppercase letters indicating differences in alkaline soil.

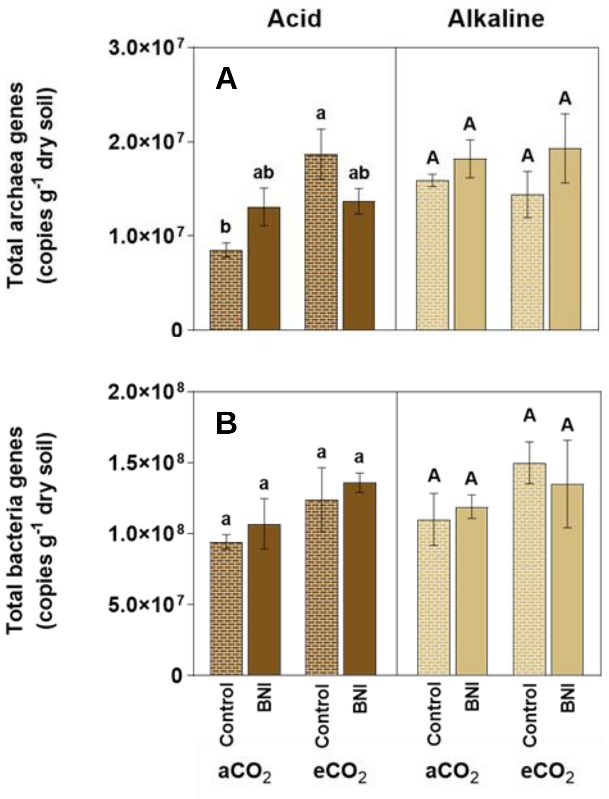


**Supplementary fig. 3.** Total abundance of (A) archaea and (B) bacteria (both measured as the abundance of their respective *16S rRNA* gene) in acidic and alkaline soils under ambient ($aCO_2$) and elevated ($eCO_2$) $CO_2$ conditions. Duncan's post hoc test ($p < 0.05$) was used to denote significant differences with lowercase letters indicating differences in acidic soil and uppercase letters indicating differences in alkaline soil.

**Supplementary table 2.** Dry aboveground biomass of ROELFS-BNI plants in acidic and alkaline soils under ambient ($aCO_2$) and elevated ($eCO_2$) $CO_2$ conditions after 45 days of growth. Duncan's post hoc test ($p < 0.05$) was used to denote significant differences with lowercase letters indicating differences in acidic soil and uppercase letters indicating differences in alkaline soil.

| | | | Aboveground biomass ($g\ DW\ plant^{-1}$) |
|---|---|---|---|
| **Acidic** | **$aCO_2$** | **Control** | 3.0 ± 0.05 b |
| | | **BNI** | 3.1 ± 0.12 b |
| | **$eCO_2$** | **Control** | 3.5 ± 0.01 a |
| | | **BNI** | 3.6 ± 0.05 a |
| **Alkaline** | **$aCO_2$** | **Control** | 3.3 ± 0.05 B |
| | | **BNI** | 3.2 ± 0.07 B |
| | **$eCO_2$** | **Control** | 3.6 ± 0.04 A |
| | | **BNI** | 3.5 ± 0.10 AB |

**Supplementary table X.** Statistical analysis of the results was made through analysis of variance (two-way ANOVA) showing the effect of $CO_2$ (C), presence/absence of BNI trait (B), and their interaction (CxB). Significant differences are marked with an asterisk (*) when $p < 0.05$, double asterisk (**) when $p < 0.01$, and triple asterisk (***) when $p < 0.001$.

| | **Acidic** | | | | **Alkaline** | | |
|---|---|---|---|---|---|---|---|
| | **C** | **B** | **CxB** | | **C** | **B** | **CxB** |
| | ** | *** | * | Accumulated $N_2O$-N emissions | ** | ** | NS |
| | *** | NS | NS | Accumulated $CH_4$-C emissions | ** | NS | NS |
| | NS | * | NS | AOA *amoA* genes | *** | *** | * |
| | NS | ** | NS | AOB *amoA* genes | ** | * | * |
| | NS | * | NS | Archaeal Chao1 | NS | * | NS |
| | NS | NS | NS | Archaeal Shannon | NS | * | NS |
| | *** | NS | NS | Bacterial Chao1 | * | NS | NS |
| | *** | NS | NS | Bacteria Shannon | NS | * | NS |
| | NS | NS | ** | *nirK* genes | * | ** | NS |
| | NS | NS | NS | *nirS* genes | * | NS | NS |
| | * | * | NS | *nosZI* genes | ** | *** | ** |
| | NS | NS | NS | *nosZII* genes | NS | NS | NS |
| | *** | NS | NS | Dry aboveground biomass | ** | NS | NS |
| | ** | NS | NS | Soil $NH_4^+$-N content | NS | NS | * |
| | * | NS | NS | Soil $NO_3^-$-N content | NS | * | * |
| | * | NS | * | Total archaea genes | NS | NS | NS |
| | NS | NS | NS | Total bacteria genes | NS | NS | NS |
| **Archaea Class** | *** | NS | * | Bathyarchaeia | NS | NS | NS |
| | ** | ** | ** | Methanobacteria | * | NS | * |
| | NS | NS | NS | Methanocellia | NS | ** | NS |
| | * | NS | NS | Methanosarcinia | NS | NS | NS |
| | NS | * | * | Nanoarchaeia | NS | NS | NS |
| | NS | * | NS | Nitrososphaeria | NS | * | NS |
| | NS | * | NS | Thermoplasmata | NS | * | NS |
| **Nitrifying archaea** | NS | NS | NS | Group_1.1c | NS | NS | NS |
| | NS | ** | NS | Nitrososphaeraceae | NS | ** | NS |
| | NS | NS | NS | Nitrosotaleaceae | NS | NS | NS |
| **Nitrifying bacteria** | NS | NS | NS | Beijerinckiaceae | NS | NS | * |
| | * | ** | * | Mycobacteriaceae | * | NS | NS |
| | NS | ** | NS | Nitrosomonadaceae | NS | ** | NS |
| | NS | ** | NS | Nitrospiraceae | NS | ** | NS |
| | NS | NS | NS | Archangiaceae | NS | NS | NS |
| | NS | NS | NS | Desulfarculaceae | NS | NS | NS |
| | NS | NS | NS | Geobacteraceae | NS | NS | NS |
| | NS | NS | NS | Halieaceae | NS | NS | NS |
| | * | NS | NS | Isosphaeraceae | NS | NS | NS |
| | NS | NS | NS | Phycisphaeraceae | NS | NS | NS |
| | NS | NS | NS | Pirellulaceae | * | NS | NS |
| | NS | NS | NS | Polyangiaceae | NS | NS | NS |
| **NO producers** | NS | NS | NS | Acidothermaceae | NS | NS | NS |
| | NS | NS | NS | Anaerolineaceae | * | NS | * |
| | NS | NS | NS | Chloroflexaceae | NS | NS | NS |
| | ** | NS | NS | Gemmatimonadaceae | NS | NS | NS |
| | NS | NS | NS | Geodermatophilaceae | NS | NS | NS |
| | NS | NS | NS | Herpetosiphonaceae | NS | NS | NS |
| | NS | NS | NS | Iamiaceae | NS | NS | NS |
| | NS | NS | NS | Micromonosporaceae | * | ** | NS |
| | NS | NS | NS | Opitutaceae | NS | NS | NS |

| | | | | | | | |
|---|---|---|---|---|---|---|---|
| | NS | NS | NS | Pseudonocardiaceae | NS | NS | NS |
| | NS | NS | NS | Roseiflexaceae | NS | NS | NS |
| | ** | NS | NS | Verrucomicrobiaceae | NS | NS | NS |
| **$N_2O$ producers** | NS | NS | NS | Cellvibrionaceae | NS | NS | NS |
| | NS | NS | NS | Chitinophagaceae | * | NS | NS |
| | NS | NS | NS | Gallionellaceae | NS | NS | NS |
| | NS | NS | NS | Methylacidiphilaceae | NS | NS | NS |
| **$N_2$ producers** | NS | NS | NS | Burkholderiaceae | NS | ** | NS |
| | NS | * | ** | Flavobacteriaceae | NS | * | NS |
| | NS | NS | NS | Hymenobacteraceae | NS | NS | NS |
| | NS | NS | NS | Pseudomonadaceae | NS | NS | NS |
| | NS | ** | NS | Rhodanobacteraceae | NS | * | NS |
| | NS | NS | NS | Rhodocyclaceae | NS | NS | NS |
| | NS | NS | NS | Sphingobacteriaceae | NS | NS | NS |
